# Does novelty influence foraging decision of a scavenger?

**DOI:** 10.1101/2023.12.12.571372

**Authors:** Debottam Bhattacharjee, Shubhra Sau, Jayjit Das, Anindita Bhadra

## Abstract

Acquiring knowledge about the environment is crucial for survival. Animals, often driven by their explorative tendencies, gather valuable information regarding food resources, shelter, mating partners, etc. While neophilia, or the preference for novel environmental stimuli or objects, can promote exploratory behaviour, neophobia, or avoidance of novel environmental stimuli, can constrain such behaviour. Neophobia can reduce predation risk, yet decreased exploratory behaviour resulting from it may limit the ability to discover potentially highly rewarding resources. Dogs (*Canis familiaris*) living in semi-urban and urban environments as free-ranging populations are, although subject to various selection forces, typically have negligible predation pressure. These dogs are scavengers in human-dominated environments; thus, selection against object-neophobia can provide benefits when searching for novel food resources. Although pet and captive pack-living dogs are known to exhibit neophilia when provided with novel objects, little is known about free-ranging dogs’ behavioural responses to novel objects, particularly in foraging contexts. Using an object choice experiment, we tested 274 free-ranging dogs from two age classes, adult and juvenile, to investigate their object-neophobia in a scavenging context. We employed a between-subject study design, providing dogs with a familiar and a potentially novel object, both baited with equal but (partially)hidden food rewards. Adult and juvenile dogs significantly inspected the novel object first compared to the familiar one, even when we reduced the visual obscurity of the familiar object, i.e., when the hidden food reward was partially visible. Furthermore, novel objects with varying strengths of olfactory cues (baited vs. false-baited) were inspected comparably by adults and juveniles. No significant differences were found in the latencies to inspect the objects. These results indicate that free-ranging dogs, evidently from an early ontogenic phase, do not show object-neophobia, as demonstrated by their preference for novel over familiar food sources. We conclude that little to no constraint of neophobia on exploratory behaviour, yet selection for object-neophilia in semi-urban and urban-dwelling animals can guide foraging decision-making processes, providing adaptive benefits.

## Introduction

Knowledge about the immediate environment is essential for survival. Animals navigate their surroundings and gather crucial information on food sources, mating partners, shelter, and predators (Mettke-Hofmann, Winkler & Leisler, 2002; Dall et al., 2005; Moretti et al., 2015; Sarkar & Bhadra, 2022), often driven by their exploratory behaviour. Exploration may further lead to innovative problem-solving (Wat, Banks & McArthur, 2020; Klump et al., 2022). Thus, exploratory behaviour directly or indirectly influences the survival and reproduction of animals. However, neophobia, or avoiding an object or other environmental aspects solely because it has never been experienced, can constrain exploratory behaviour (Stöwe et al., 2006a). While a suppressed tendency to explore can decrease the risk of encountering predators, it can also substantially limit opportunities to discover novel resources, like food, mating partners, or shelters (Stöwe et al., 2006b). In contrast to neophobia, neophilia is a trait which can promote exploratory behaviour, where animals prefer novelty over familiarity (Day et al., 2003). Notably, neophobia and neophilia are thought to be shaped by different selective factors; thus, they do not necessarily represent two extremes of a continuum (Greenberg & Mettke-hofmann, 2001). Furthermore, neophobic and neophilic responses can be highly context-specific, with studies typically using neophilia in neutral and neophobia in foraging contexts, respectively (Takola et al., 2021). Nevertheless, a complex interaction between the two can exist, e.g., individuals (or species) explore (i.e., driven by neophilia) for survival but may do so with high levels of fear and arousal (i.e., neophobia) to keep themselves prepared for any potential adversities associated with novelty (Greenberg, 2003). Animals inhabiting urban and semi-urban (i.e., human-dominated) environments experience a lower predation rate than those in rural and wild habitats (Fischer et al., 2012; Eötvös, Magura & Lövei, 2018). Therefore, suppressed object-neophobia in urban-dwelling animals can enhance exploratory behaviour, providing benefits. A long-standing bias of scientists, especially behavioural ecologists, was to ignore ‘subsidised’ animals that ‘exploit’ anthropogenic food resources. Consequently, our knowledge of these animals’ neophilia/neophobia-related behavioural responses is obscured. Nevertheless, with the establishment of urban ecology, scientific research has shifted gears towards understanding the behavioural and cognitive aspects of decision-making of animals living close to humans.

Human-dominated environments impose novel challenges on animals (Ditchkoff, Saalfeld & Gibson, 2006). Object-neophilia (and a reduced object-neophobia) and exploratory behaviour are crucial behavioural adaptations to thrive in these environments (Griffin, Netto & Peneaux, 2017), which are often repeatable (i.e., personality) and heritable (Day et al., 2003; Quinn et al., 2009). For instance, species that invade human-dominated environments are known to be neophilic (Sol, Lapiedra & González-Lagos, 2013). Several studies suggest that urban-living animals have reduced object-neophobia (Tryjanowski et al., 2016; Greggor et al., 2016; Jarjour et al., 2019; Biondi et al., 2020; Miller et al., 2022), but see (Mazza et al., 2021). Yet, plasticity in neophobia can be seen with higher human disturbance levels, and shifts in personality types are assumed in human-dominated environments, involving reductions in neophobia (Grunst et al., 2019). Furthermore, empirical studies provide evidence that a wide range of species living close to humans exhibit enhanced exploratory behaviour (Thompson et al., 2018; Breck et al., 2019; Dammhahn et al., 2020), and the underlying proximate mechanisms that regulate exploration are based on neophobia-neophilia related traits (Griffin, Netto & Peneaux, 2017). Despite a wealth of knowledge gathered by empirical studies on those traits, our understanding of the ecologically relevant contexts under which such traits can be expressed and/or be plastic is lacking (see Gordon, 2011). Consequently, testing more species that live in close proximity to humans in ecologically relevant contexts, such as foraging, is imperative.

Dogs (*Canis familiaris*) inhabit a wide range of habitats, e.g., living as pets in human households or as free-ranging populations in human-dominated environments. Unlike pets, free-ranging dogs are primarily under natural and sexual selection pressures (Range & Marshall-Pescini, 2022); thus, acquiring information about their surroundings is key to their survival. However, little is known about whether these dogs exhibit object-neophobia. Several studies highlight free-ranging dogs’ socio-cognitive abilities, potentially guiding their decision-making processes and making them successful in human-dominated environments. Similar to other ‘urban adapters’ or ‘urban exploiters’ (McKinney, 2006), these dogs maintain a wary distance from humans but can build trust with or learn socially from unfamiliar humans (Bhattacharjee et al., 2017d; Cimarelli et al., 2023). Moreover, their abilities to understand human communicative intents, including gestures, attentional states and facial expressions, have recently been evidenced by empirical studies (Bhattacharjee et al., 2019; Brubaker et al., 2019; Bhattacharjee & Bhadra, 2022; Lazzaroni et al., 2023). Free-ranging dogs also show a high degree of behavioural plasticity and engage with various objects provided by humans to perform tasks that lead to food rewards (Bhattacharjee et al., 2017c, 2019; Lazzaroni et al., 2019; Cimarelli et al., 2023), often using judgment to choose the best available option (Bhattacharjee et al., 2017c, 2019). Yet, as scavengers, how non-social traits, such as neophobia and/or neophilia, shape free-ranging dogs’ decision-making processes has rarely been investigated.

Exploratory behaviour in free-ranging dogs can be of significant adaptive value. They spend considerable time and energy walking and scavenging (Sen Majumder, Chatterjee & Bhadra, 2014), which may include exploring the surroundings and potentially encountering novel challenges or objects, such as human artefacts. Free-ranging dogs exhibit varying scavenging strategies, from following simple rules of thumb for foraging decisions to extractive foraging techniques (Mangalam & Singh, 2013; Bhadra et al., 2015). However, it is unknown whether, in general, such behaviours are influenced by object-neophobia. Solitary foraging is prevalent in these dogs (Sen Majumder et al., 2014) to reduce intraspecific competition (Sarkar, Sau & Bhadra, 2019; Sarkar et al., 2023). At the same time, these dogs do not have any natural predators in human-dominated environments, where humans are the biggest cause of their mortality (Paul et al., 2016). In the urban environment, they are likely to encounter novel human artefacts, some of which might be rewarding if explored, like garbage bins, packaged food and containers (personal observation). Thus, individuals may benefit from the lack of object-neophobia. Pet and captive pack-living dogs exhibit little to no neophobia and enhanced neophilia (Kaulfuß & Mills, 2008; Moretti et al., 2015) in choice tasks when provided with familiar and unfamiliar objects. The presence of group members in captive pack-living dogs further induces exploration, indicating risk-sharing (Moretti et al., 2015). However, considering early age classes, a dog-wolf comparative study suggests that wolves are more persistent in exploring a novel environment and novel objects when compared with dogs (Marshall-Pescini et al., 2017b). Nevertheless, to our knowledge, no study investigated whether free-ranging dogs exhibit object-neophobia, particularly in scavenging contexts and if such a trait influences their foraging decisions.

Here, we conducted a simple yet ecologically relevant object-choice test with free-ranging dogs to investigate whether their foraging decisions depend on (or are constrained by) object-neophobia. A between-subject study design was employed, and dogs were provided with one of the following combinations of objects – (a) an opaque plastic ball and an opaque plastic pouch (test), (b) an opaque plastic ball and a translucent plastic pouch (test), (c) an opaque plastic pouch and a translucent plastic pouch (control), and (d) two opaque plastic balls (validation). All the objects contained equal-sized food rewards, except for the last condition, where one of the balls was false-baited by gently rubbing a piece of food reward. Therefore, the first three conditions (a, b, and c) had varying levels of visual obscurities but similar olfactory cues, but the last condition had only different strengths of olfactory cues. Freeranging dogs typically encounter plastic garbage bags and pouches and extract food leftovers from them (Bhattacharjee et al., 2017a). On the contrary, plastic balls are unlikely to be encountered during scavenging (personal observation). Accordingly, plastic balls and pouches were considered novel- and familiar objects, respectively. We hypothesised that free-ranging dogs, being scavengers in human-dominated environments, will not exhibit object-neophobia. In particular, we expected dogs to inspect the novel objects first rather than the familiar ones (in test conditions a and b). Moreover, since enhanced neophilia is considered an adaptive trait in pet dogs (Kaulfuß & Mills, 2008), we expected both adults and juveniles to respond similarly, i.e., no ontogenic differences in their behavioural responses. For the control and validation conditions (conditions c and d), we expected dogs to exhibit comparable choices, i.e., no clear preference between the objects. Finally, we predicted that dogs, if they are not neophobic, will show comparable latencies in approaching the objects under different conditions.

## Methods

### (i) Ethical statement

In India, free-ranging dogs are protected by the Prevention of Cruelty to Animals Act (1960) of Parliament, which allows interactions with dogs, including feeding and petting. Our study adhered to the guidelines of the act and to the ethical guidelines of animal testing of the Indian Institute of Science Education and Research Kolkata (Approval no. 1385/ac/10/CPCSEA). The subject dogs were tested in their natural habitats, and their participation in the tests was voluntary. All meat items used were fresh and fit for human consumption.

### (ii) Study area and subjects

The study sites were the following semi-urban and urban areas of West Bengal, India - Mohanpur (semi-urban, 22°56’49’’N and 88°32’4’’E), Kalyani (semiurban to urban, 22°58’30” N, 88°26’04” E) and Kolkata (urban, 22°57’26” N, 88°36’39” E). We covered a total sampling area of 88 Km^2^ (Mohanpur: 11 Km^2^, Kalyani: 21 Km^2^, and Kolkata: 56 Km^2^). The study was conducted from October 2016 to May 2017, between 0900 and 1800 hours. The experimenters walked on the streets to locate dogs, preferably present without group members. In case more than one dog was present, a focal subject dog was chosen randomly and lured out of sight of the group members.

We tested 274 free-ranging dogs from two age classes: adults (n=147) and juveniles (n=127). While the exact age of the dogs was unknown, morphological characteristics enabled us to define an age class precisely (Sen Majumder et al., 2014). The sexes of the dogs were noted by visually inspecting their genitalia. Of the 147 adult dogs, 77 were females, whereas 63 out of 127 were females in the juvenile age class. Thus, the male-female ratio of our overall sample was close to 1:1. Since tracking a large sample of dogs over the experimental period is highly challenging, we decided to use a between-subject study design. All dogs were tested once, and to rule out any potential bias of resampling, we tested dogs from different locations (i.e., the large 88 Km^2^ sampling area) and never revisited the same area for testing. Notably, the participation of the dogs in the study was voluntary.

### (iii) Experimental objects

We used three different objects in this study - a green opaque plastic ball with a diameter of 2.8″, a black opaque plastic pouch of 7.5″ × 4.5″ size, and an identically sized white translucent plastic pouch. For each experimental trial, new and completely unused objects were used.

### (iv) Experimental conditions and procedure

Upon locating an adult or a juvenile dog, the experimenter presented them with one of the following experimental conditions randomly – (a) a green opaque plastic ball and a black opaque plastic pouch (GB-BP; **Video S1**), (b) a green opaque plastic ball and a translucent white plastic pouch (GB-TP; **Video S2**), (c) a black opaque plastic pouch and a translucent white plastic pouch (BP-TP; **Video S3**), and (d) two green opaque plastic balls (GB-GBB; **Video S4**) (see **Table 1** for descriptions). We used raw chicken pieces (∼ 15 g) as food items. A circumferential opening was made in the plastic ball to place the food item. Later, the opening was loosely closed using transparent tape such that the structure of the ball remained intact. Similarly, after placing the food items inside, the openings of the plastic pouches were loosely tied with cotton threads. Each object from the GB-BP, GB-TP, and BP-TP conditions contained a food item inside. In the GB-GBB condition, one of the plastic balls (GB) had a food item similar to previous conditions, but the other ball (GBB) was false-baited by gently rubbing a food item inside. Thus, despite appearing identical, GB had a food item inside while GBB was empty, thus providing varying strengths of olfactory cues. In summary, the experimental conditions provided similar visual and olfactory cues (GB-BP), similar olfactory yet different visual cues (GB-TP, BP-TP), and similar visual with different strengths of olfactory cues (GBB-GB) to the dogs.

**Table 1:** The experimental conditions and their details. A tabular summary of the experimental conditions, corresponding objects used, and condition details.

| Experimental condition | Object | Description |
| --- | --- | --- |
| (a) GB-BP | GB | Familiar (BP) and novel (GB) objects with similar visual obscurity (opacity) and olfactory cues. Test condition to check which object is preferred. |
|  | BP |  |
| (b) GB-TP | GB | Familiar (TP) and novel (GB) objects with different visual obscurity but similar olfactory cues. Test condition to check which object is preferred when the familiar object has partial visual cues of the reward. |
|  | TP |  |
| (c) BP-TP (Control) | BP | Familiar objects with varying visual obscurity levels but similar olfactory cues. Control condition |
|  | TP | to check if they are perceived similarly as familiar objects or discriminated against based on visual cues. |
| (d) GB-GBB (Validation) | GB | Novel objects with similar visual obscurity but varying strengths of olfactory cues. GB and GBB with and without food items inside, respectively. GBB is false-baited by gently rubbing the food reward inside, thus providing a relatively lower strength of olfactory cue than GB. Validation condition to check if olfactory cues determine object choice over novelty. |
|  | GBB |  |

The abovementioned combination of the two objects was placed on the ground approximately 1 meter from each other. We used a pseudorandomised order to place the objects (left/right) to avoid any potential effects of side bias. The objects were equidistant from a focal dog. The minimum distance between the midpoint of objects and the focal dog was approximately 2 m. The experimenter stood 0.5 m behind the midpoint of the two objects and tried to get the attention of a focal dog by calling “aye aye” (Bhattacharjee et al., 2017d). After capturing the dog’s attention (i.e., when the focal dog’s head was oriented towards experimental setup), the experimenter left the set-up and positioned himself at a minimum distance of 5 m away or hid behind a tree or car. This step ensured the participation of relatively shy individuals and eliminated any potential influence of human presence and subsequent begging-related behaviour exhibited by the dogs. A trial began immediately after capturing the focal dog’s attention and lasted 1 minute. If a dog was not attentive, the experimenter made another attempt after 10 sec. A maximum of two such additional attempts were made before terminating a trial. The trials were video recorded by a person (D.B.) other than the experimenter from a distance of minimum 5 m using a handheld camera. Notably, S.S. and J.D. played the roles of the experimenter, who were both males and had a similar height and physical build. Therefore, dogs’ responses were unlikely to be influenced by the two different experimenters involved in the study. Besides, the subject dogs witnessed the experimenters briefly during a one-off test.

### (iv) Data coding

We coded two behavioural variables – object choice and the latency to choose the object. S.S. coded the videos using a frame-by-frame video inspection method. Another rater coded 15% of the videos to check for reliability, and it was found excellent (Intraclass correlation coefficient/ICC test, first inspection: (ICC (3,k)) = 0.99, p<0.001; latency - (ICC (3,k)) = 0.94, p<0.001). Object choice was defined as the first physical inspection of an object by touching (with the muzzle) or licking (with the tongue). Thus, a clearly visible physical interaction between dogs and the objects was considered. We noted which object was inspected first (i.e., GB or BP, GB or TP, BP or TP, and GB or GBB) in all four conditions. Latency was defined as the time taken by a focal dog to inspect an object from an initial 2 m distance. Due to the potentially varying difficulty levels associated with food retrieval from the objects, we decided not to code object-based activities (or exploration behaviour, though in a strict sense) after the first inspection—nonetheless, object choice in itself incorporates the primary crucial step of foraging decision-making. Of the 274 dogs, 15 (nine adults and six juveniles) did not move from their initial position. Subsequently, we conducted our analyses on a revised sample of 259 free-ranging dogs.

### (v) Statistics

All statistical analyses were conducted using R version 4.3.1. (R Development Core Team, 2019). We used binomial tests for the four experimental conditions separately to check whether dogs preferred one object over another. Generalised linear models (GLM) were used to investigate if age class (adult and juvenile) influenced the decision of object choice. Four binomial GLMs were run, each for the different experimental conditions, where the type of object and age class were included as response and independent variables, respectively. In addition, we added the sexes of the dogs (male and female) as control variables in those models. We used linear effect models (LM) to investigate the latency of first inspection. Latency was included as the response variable, and age class, sex (as a control variable), and the corresponding object inspected were included as fixed effects in the models. We carried four LMs for the different experimental conditions. GLM and LM were conducted using the “lme4” package (Bates et al., 2015). In case full models had significant effects, comparisons with null models were checked using the “lmtest” package (Hothorn & Zeileis, 2011). Model diagnostics were checked using the “DHARMa” package (Hartig, 2020). If model residuals violated normality assumptions, we log-transformed the response variable and re-ran the model. The level of statistical significance (α) was set at 0.05.

## Results

### (i) First inspection (object choice)

a. *GB-BP condition –* 62% (23 out of 37) and 69% (18 out of 26) of the adults and juveniles inspected GB first, respectively. Regardless of age classes, dogs first inspected GB significantly more over BP (Binomial test: p = 0.02, 95% Confidence Interval/CI = 0.52, 0.76, **Figure 1**). However, we found no significant effect of age class (GLM: z = 0.6, p = 0.54) and sex (GLM: z = 0.3, p = 0.76) on object choice. These results suggest that both adults and juveniles comparably preferred the novel object over the familiar one.

**Figure 1:**
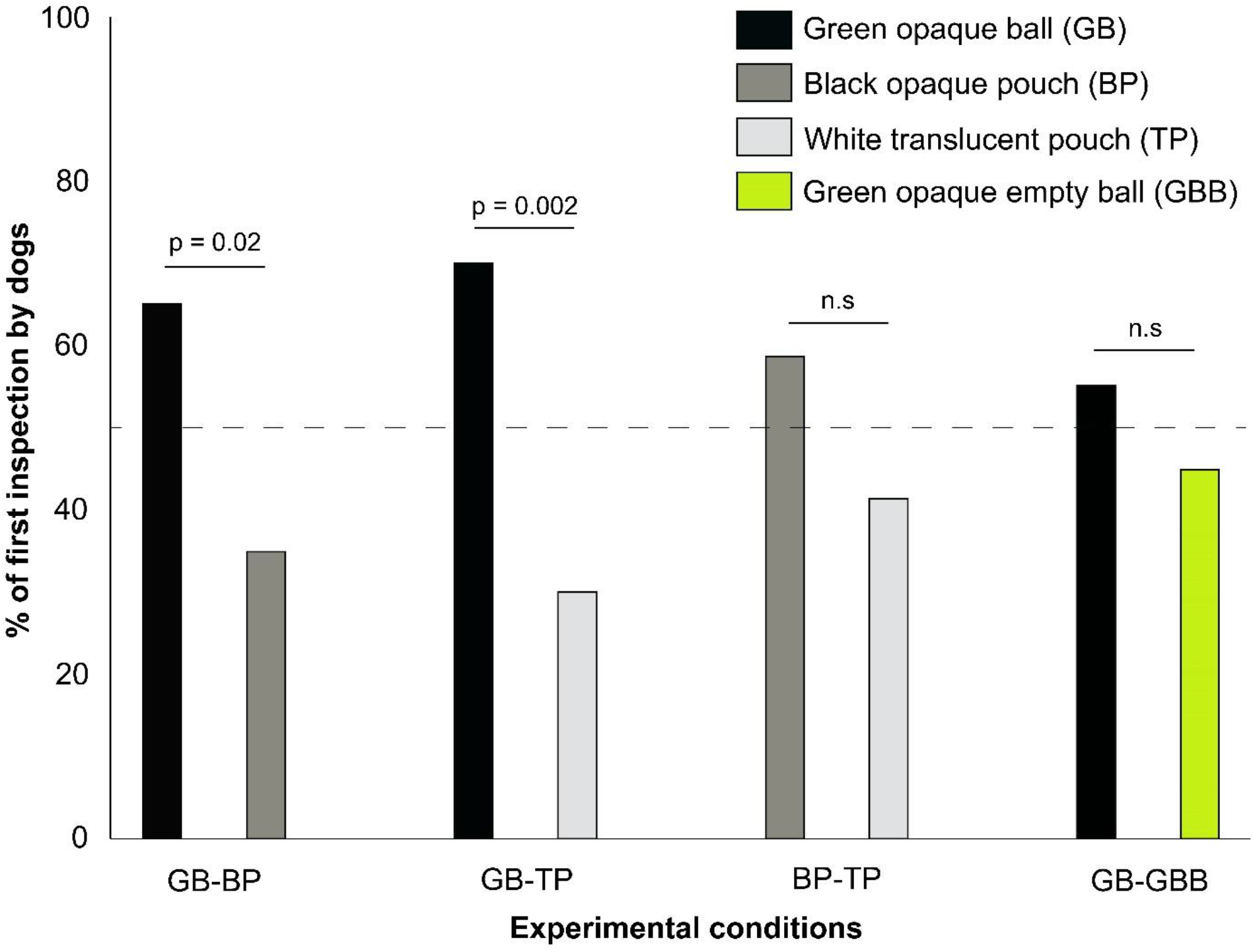
A bar graph showing the percentage of first inspection by dogs in different conditions. Significant differences are highlighted with p values and non-significant differences are denoted by “n.s”. GB-BP and GB-TP represent test conditions, whereas BP-TP and GB-GBB represent control and validation conditions, respectively.
b. *GB-TP condition –* 73% (25 out of 34) and 65% (17 out of 26) of the adults and juveniles inspected GB first, respectively. Overall, dogs first inspected GB significantly more than TP (Binomial test: p = 0.002, 95% CI = 0.57, 0.81, **Figure 1**). Age class (GLM: z = -0.68, p = 0.49) and sex (GLM: z = 1.12, p = 0.26) did not influence object choice. Thus, similar to the GB-BP condition, adults and juveniles first inspected the novel object more than the familiar object, even when the familiar object had visual cues.
c. *BP-TP condition –* 52% (15 out of 29) and 63% (19 out of 30) of the adults and juveniles inspected BP first over TP, respectively. In general, dogs did not differ in their first inspection of the familiar objects (Binomial test: p = 0.29, 95% CI = 0.44, 0.70, **Figure 1**), even when they differed in visual obscurity levels. No significant effect of age class (GLM: z = 0.78, p = 0.43) and sex (GLM: z = -0.5, p = 0.61) on object choice was found.
d. *GB-GBB condition –* 56% (22 out of 39) and 54% (21 out of 39) of the adults and juveniles first inspected GB, respectively. Overall, dogs did not discriminate between the novel objects during the first inspection, even when the strengths of olfactory cues differed (Binomial test: p = 0.49, 95% CI = 0.42, 0.65, **Figure 1**). We did not find any significant effect of age class (GLM: z = -0.33, p = 0.74) and sex (GLM: z = -0.5, p = 0.61) on object choice.

### (ii) Latency of first inspection

a. *GB-BP condition –* The latencies to inspect GB (mean ± standard deviation: 2.90 ± 3.57 seconds) and BP (2.13 ± 1.93 seconds) did not differ (LM: t = 1.27, p = 0.20). We found no effect of age class (LM: t = -1.83, p = 0.07) and sex (t = -1.28, p = 0.20) on latencies.
b. *GB-TP condition –* Dogs inspected GB (2.42 ± 1.51 seconds) comparably to TP (4.05 ± 6.43 seconds) (LM: t = 0.91, p = 0.37). No effect of age class (t = -1.13, p = 0.26) and sex (t = 0.13, p = 0.89) on the latencies was found.
c. *BP-TP condition –* The latencies to inspect BP (2.11 ± 1.57 seconds) and TP (3.16 ± 4.32 seconds) were similar (LM: t = 1.20, p = 0.23). Age class (t = 1.48, p = 0.14) and sex (t = 1.33, p = 0.18) did not impact the latencies of first inspection.
d. *GB-GBB condition –* We did not find any difference in dogs’ latencies to inspect GB (2.67 ± 2.17 seconds) and GBB (2.31 ± 1.15 seconds) (LM: t = -1.10, p = 0.27). No effect of age class (t = 0.09, p = 0.93) and sex (t = -1.15, p = 0.25) was found.

## Discussion

In this study, we investigated whether free-ranging dogs exhibit object-neophobia and whether their foraging decision-making is dependent on, and in particular, constrained by, neophobic behaviour. We tested dogs from two age classes to evaluate if such behavioural responses have any ontogenic developmental basis. As hypothesised, we found that free-ranging dogs are not neophobic, particularly in a scavenging context. Subsequently, their foraging decisionmaking, albeit at the primary yet crucial stage, is not constrained by object-neophobia. In line with our expectations, juvenile and adult dogs exhibited similar behavioural responses, indicating no ontogenic differences in neophobia in dogs. This could potentially translate into the adaptive benefits of suppressed object-neophobia in dogs and/or selection for neophilia, as previously suggested by studies on other populations of dogs (Kaulfuß & Mills, 2008; Moretti et al., 2015). Our results not only strengthen the view that dogs are not neophobic but also provide empirical evidence by designing a task which mimics an ecologically relevant context of scavenging. We discuss how reduced object-neophobia (and potentially heightened neophilia) can be crucial in free-ranging dogs present in human-dominated environments.

Human-dominated environments offer food subsidies, and these predictable resources can have broad ecological and evolutionary implications for animals (Oro et al., 2013). Despite their predictability, a wide range of species exploit these resources (Biswas et al., 2022), resulting in substantial competition both within and between species. Furthermore, anthropogenic activities can alter the spatiotemporal availability of those resources and directly or indirectly determine how animals eventually utilise them (Markus & Hall, 2004; Murray & St. Clair, 2017; Ramírez et al., 2020; Bhattacharjee & Bhadra, 2021; Egert-Berg et al., 2021). To maximise their food intake while avoiding potential human-related stressors, animals may engage in opportunistic scavenging and explore novel resources. In line with this, a reduction in object-neophobia in free-ranging dogs can, therefore, be an adaptive strategy to promote exploratory behaviour, especially during foraging.

In both the test conditions, dogs preferred the novel over the familiar objects, suggesting no direct constraint of object-neophobia on their potential exploratory behaviour. Interestingly, even when the hidden food reward was partially visible from the familiar object, dogs preferred the opaque novel object over it. This is attributed to dogs’ reliance on a complex multimodal sensory information system of vision and olfaction during foraging. Dogs are primarily known to rely on olfactory cues to make their foraging decisions (Bhadra et al., 2015; Banerjee & Bhadra, 2019; Sarkar et al., 2023), but they ‘seem to lose their nose’ in a communicative context with humans, where they rely more on visual than olfactory cues (Szetei et al., 2003). Although our study did not involve an active presence of human experimenters, particular interest towards human artefacts should be tolerated due to selection for neophilia (Kaulfuß & Mills, 2008). Contrary to the test conditions, dogs in our study did not exhibit a particular preference for familiar objects with different visual cues (BP-TP) or novel objects with different olfactory cues (GB-GBB). This might suggest that familiar food resources are of comparable values regardless of their appearance, albeit with similar olfactory cues. The GB-GBB condition, on the other hand, validates our results by demonstrating that novelty indeed played a role, and dogs did not fully rely on (the strength of) olfactory cues. These findings indicate that dogs can use their multimodal sensory information system rather flexibly, depending on the context, to guide their foraging decisions. However, it is worth mentioning that olfaction is a strong inducer for scavenging, which is why we used hidden or partially hidden food rewards to motivate dogs to approach the task setup. Nevertheless, neophobia may still be relevant in contexts with (i.e., object-neophobia) or without (i.e., object-neophilia) food (Takola et al., 2021), as discussed previously. Taken together, our results corroborate previous findings that dogs are not neophobic and instead may show signs of neophilia, which in turn can promote exploratory behaviour and guide their decision-making processes.

In free-ranging dogs, different ontogenic phases hold varying degrees of significance. The juvenile phase of development (3-6 months) involves complete independence from the mothers and venturing into the immediate environment (Paul et al., 2016); developmentally, this is a crucial phase as dogs start to forage on their own and experience anthropogenic stressors. It has been shown that juvenile free-ranging dogs are reluctant to approach and follow the communicative intents of unfamiliar humans (Bhattacharjee et al., 2017c). Therefore, it can be assumed that scavenging, but not begging for food from humans, is the predominant feeding strategy in juveniles. A reduced object-neophobia can thus provide considerable foraging benefits to juveniles. Conversely, adults can rely on scavenging and begging (Sen Majumder et al., 2014; Bhadra et al., 2015; Boitani et al., 2016), and depending on the energy requirements and other external factors (such as human disturbance), they may flexibly switch between those strategies, where reduced object-neophobia would still be beneficial. Nonetheless, object-neophobia in dogs appears to be a trait that was selected against during domestication (but see Marshall-Pescini et al., 2017b).

Studies with novel object tests often consider investigating the time animals take to approach or explore the objects. Neophilic individuals are expected to approach the novelty quicker than their less-neophilic or neophobic counterparts (Moretti et al., 2015; Takola et al., 2021). One may argue that dogs in our study did not differ in their latencies to approach the novel and familiar objects. This could be attributed to the study design, where we provided a two-way object choice condition instead of separately presenting novel or familiar objects. Besides, in contexts of scavenging, dogs’ object choice can be more important than how fast they approach, as free-ranging dogs are primarily solitary foragers. This does not disregard the potential influence of competition and subsequent shorter latencies to approach novel food sources during group foraging events. Unfortunately, we could not compare dogs’ so-called explorative behaviour towards novel and familiar objects due to the potential task difficulty levels. However, selection or, in this case, the first inspection of an object highlights an important primary step in foraging decision-making. Nonetheless, it can be assumed that object-neophobia does not constrain these dogs’ foraging decision-making. In fact, freeranging dogs are known to engage with various kinds of human artefacts during scavenging-related tasks, like baskets (Sarkar, Sau & Bhadra, 2019; Sarkar et al., 2023), bowls (Bhattacharjee et al., 2017c, 2019), containers and packaged food (Bhattacharjee et al., 2017b; Banerjee & Bhadra, 2019), etc. In future, it would also be interesting to test freeranging dogs’ exploratory behaviour towards novel and familiar objects outside of the object-choice paradigm.

Finally, socio-ecological forces can drive behavioural responses in animals. The ‘social-ecology’ hypothesis suggests that feeding ecology and social organisation may act together as mechanisms to drive dogs’ interactions with environmental, particularly novel stimuli (Marshall-Pescini et al., 2017a). Our results provide support to the hypothesis that freeranging dogs are not neophobic. Yet, it would be interesting for future studies to measure consistency in such behavioural responses and further quantification of individual-level differences. Although neophobic responses can be repeatable and present as personality traits (Sih, Bell & Johnson, 2004), potential plasticity (another key mechanism driving animals to be successful in urban habitats, see (Vincze et al., 2016; Greggor et al., 2016)) in such traits should be investigated with even broader sampling areas, especially by including rural areas with low human disturbances.

## Conclusions

We tested a large sample of free-ranging dogs in an ecologically relevant scavenging context to investigate whether they exhibit object-neophobia, and if their foraging decision-making is potentially constrained by the same. This is the first experimental evidence that free-ranging dogs do not show object-neophobia. We assume that selection against object-neophobia may provide dogs with considerable foraging benefits in a human-dominated environment. Our results support the ‘socio-ecology’ hypothesis that further explains the potential driving mechanisms behind reduced neophobia in dogs. Yet we propose examining consistency and possible plasticity in such a trait to understand its proximate mechanisms and evolution better.

## Acknowledgements

We thank Dr. Susnata Karmakar for allowing us to use his vehicle during fieldwork. We thank the Indian Institute of Science Education and Research Kolkata for providing infrastructural support.

## Additional Information and Declarations

### Funding

The work was funded by a SERB grant awarded to A.B (Department of Science and Technology, Govt. of India, project no. EMR/2016/000595). D.B. was supported by a DST INSPIRE Fellowship, Department of Science and Technology, Govt. of India. J.J. was supported by a DST INSPIRE Scholarship for Higher Education, Department of Science and Technology, Govt. of India. The funders had no role in study design, data collection and analyses, decision to publish, or preparation of the manuscript.

## Grant Disclosures

SERB Grant, Department of Science and Technology, Govt. of India (Project no. EMR/2016/000595)

DST INSPIRE Fellowship, Department of Science and Technology, Govt. of India.

DST INSPIRE Scholarship for Higher Education, Department of Science and Technology, Govt. of India.

## Competing Interests

The authors declare that there are no competing interests.

## Author Contributions

- Debottam Bhattacharjee conceived and designed the experiments, performed the experiments, analysed the data, prepared figures and/or tables, authored or reviewed drafts of the paper, approved the final draft.
- Shubhra Sau performed the experiments, analysed the data, approved the final draft.
- Jayjit Das performed the experiments and approved the final draft.
- Anindita Bhadra conceived and designed the experiments, authored or reviewed drafts of the paper, approved the final draft.

## Animal Ethics

Ethical approval of the study was obtained from Indian Institute of Science Education and Research Kolkata (Approval no. 1385/ac/10/CPCSEA).

## Field Study Permissions

No permission outside of the Institute ethics committee is required to perform non-invasive studies with free-ranging dogs in India.

## Data Availability

Data and R-Script will be made available upon publication.

## Notes

### Competing Interest Statement

The authors have declared no competing interest.

